# A bacterial endosymbiont with a highly reduced genome forms extensive membrane contact sites with host organelles

**DOI:** 10.64898/2026.10.06.757092

**Authors:** Dalton J. Leprich, Jesse Aaron, Eleni Hughes, Mark S. Ladinsky, Pamela J. Bjorkman, John P. McCutcheon

## Abstract

Bacterial endosymbionts that establish long-term associations inside of host cells often lose a huge number of genes and become dependent on their hosts for essential cellular functions^1,2^. One such example is the mealybug endosymbiont *Tremblaya princeps*: its ~139-kb genome encodes only ~120 protein coding genes and lacks all recognizable genes for lipid and fatty acid synthesis, cell wall components, membrane transport, protein translocation, and transmembrane proteins, raising fundamental questions of how Tremblaya’s cellular envelope is formed and maintained^3,4^. Here, using a combination of transmission electron microscopy (TEM) and focused ion beam scanning electron microscopy (FIB-SEM) of an entire endosymbiont-containing insect cell, we identified extensive and specific contacts between host cell mitochondria and *Tremblaya*. These contacts occurred at close membrane appositions and contained electron-dense bridge-like structures closely resembling classical eukaryotic membrane contact sites. Whole-cell segmentation showed that approximately 53% of host mitochondria were in contact with *Tremblaya*, with some mitochondria extensively wrapped by *Tremblaya*-associated membranes. Mitochondria in contact with *Tremblaya* were larger and contained proportionally greater cristae volume and surface density compared to non-contact mitochondria, exhibiting differences in both external and internal mitochondrial architecture. As membrane contact sites are well known to mediate lipid and metabolite exchange between membranes^5–10^, these findings suggest that direct interactions with host organelles may be a source of vital membrane and cellular components for endosymbionts that have lost the autonomous ability to produce and transport these molecules.

## RESULTS

### The origins and host interactions of Tremblaya’s three membranes

The mealybug *Planococcus citri* contains nested bacterial endosymbionts, where one endosymbiont, *Candidatus* Moranella endobia (hereafter *Moranella*), resides in the cytoplasm of another bacterial endosymbiont, *Candidatus* Tremblaya princeps (hereafter *Tremblaya*), which itself resides in the cytoplasm of specialized host cells called bacteriocytes (**Figure 1A**)^11,12^. In adult insects, *Tremblaya* is surrounded by three membranes (**Figure S1B, S1F**). We suspected that the outermost membrane was a host-derived symbiosomal membrane, produced from phagocytosis of *Tremblaya* cells by bacteriocytes during embryo development. We tested this idea using TEM to image *Tremblaya* cells in the extracellular “symbiont ball” of mealybug eggs, before the endosymbionts were phagocytosed into insect host cells, and verified that *Tremblaya* at this stage retained only two membranes, not three (**Figure S1C-E, S1G**). We infer that the third, outermost membrane is lost during exocytosis to eggs and regained during phagocytosis during embryogenesis^13^. We can therefore divide the three membranes of *Tremblaya* into two ancestral sources: the outermost membrane is host-derived, originating from phagocytosis and likely a modified phagosomal membrane (aka, symbiosomal membrane) similar to that reported in aphids^14,15^, and the inner two membranes are likely remnants of the diderm structure of the betaproteobacterial ancestry of *Tremblaya*^16^. However, the origin of these two inner membranes remains obscure as *Tremblaya* retains no genes to synthesize these structures^4^.

**Figure 1:**
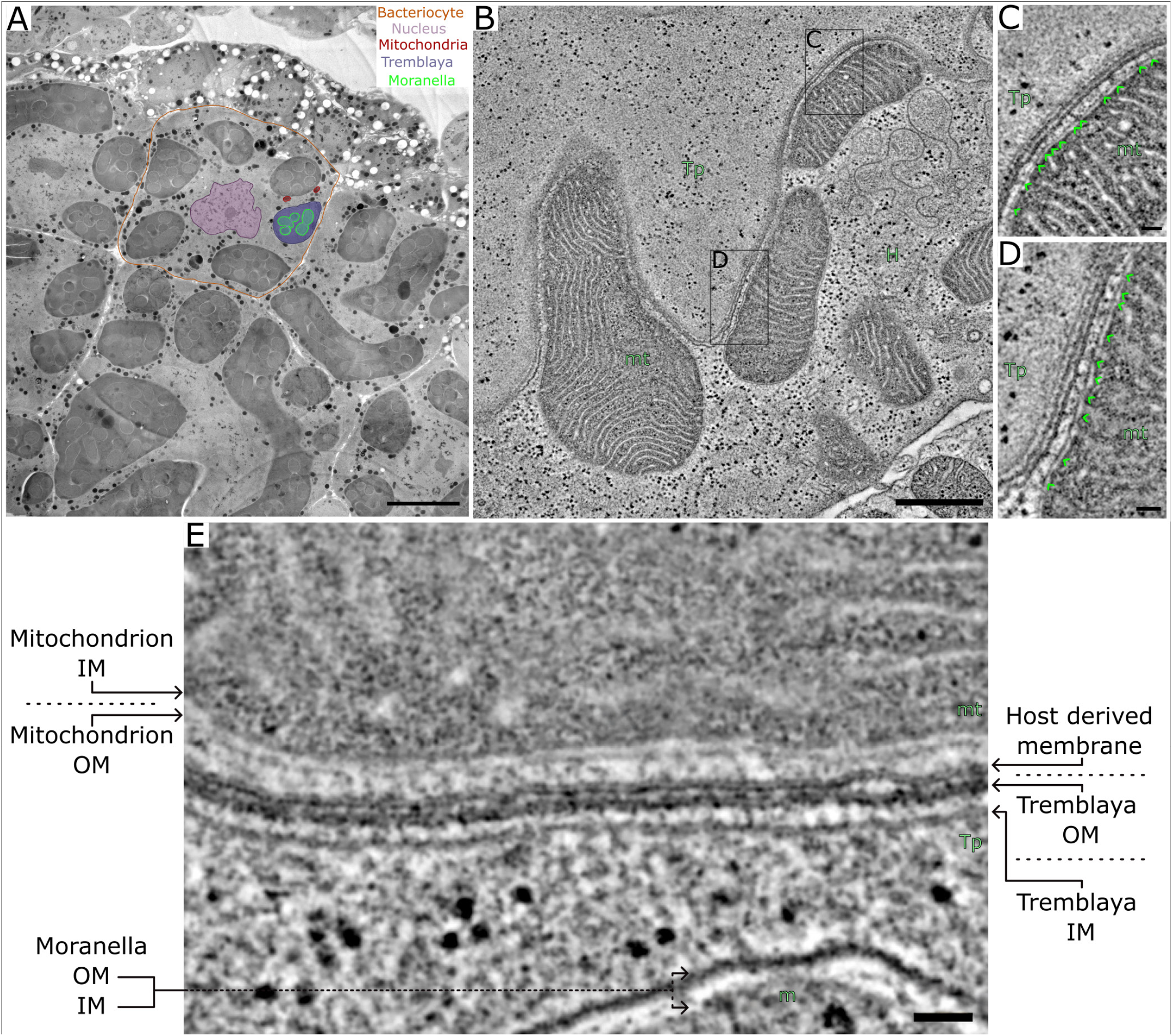
Electron-dense connections between host mitochondria and the *Tremblaya*-associated symbiosomal membrane. (A) Low-magnification TEM micrograph of a *P. citri* bacteriome. A representative bacteriocyte is outlined in orange, along with the host nucleus, mitochondria, *Tremblaya*, and *Moranella* highlighted as indicated. (B) Tomogram slice showing mitochondria positioned in close proximity to *Tremblaya*. Boxed regions are enlarged in (C) and (D). (C, D) Higher-magnification TEM views of mitochondria-*Tremblaya* interfaces. Green arrowheads indicate electron-dense, bridge-like tethers spanning the region between the outer mitochondrial membrane and the host-derived symbiosomal membrane surrounding *Tremblaya*. (E) High magnification TEM micrograph of a mitochondrion in close apposition to *Tremblaya*. Abbreviations: Tp, *Tremblaya*; m, *Moranella*; H, host cell, mt, mitochondrion. Scale bars: A, 10 µm; B, 0.5 µm; C-E, 0.05 µm.

Because *Tremblaya* lacks recognizable genes for lipid synthesis, membrane transport, and protein translocation, we examined whether its membranes might interact directly with host organelles. TEM and electron tomography on high-pressure frozen bacteriomes revealed close membrane appositions between host mitochondria and *Tremblaya* (**Figure 1B**). At these interfaces, electron-dense bridge-like structures spanned the gap between the outer mitochondrial membrane and the symbiosomal membrane (**Figure 1C-E**). These structures resemble the tethered architecture of established organelle membrane contact sites (MCS), which are structures known for lipid synthesis and exchange^5,8,10,17,18^. The presence of MCSs between *Tremblaya* and host mitochondria suggested a potential lipid acquisition mechanism for *Tremblaya*, although the extent of these MCSs remained unclear because the thin tissue sections required by TEM prevented us from quantifying the extant of these structures in whole bacteriocyte tissue.

Tomographic reconstruction of TEM semi-thick sections revealed extensions emanating from the host-derived symbiosomal membrane surrounding *Tremblaya*, and to some degree the ancestral diderm membranes, into the surrounding bacteriocyte cytoplasm (**Figure S2**). Tomographic reconstruction revealed these tentacle-like projections (TLPs) remained continuous with the symbiosomal membrane and were found extending hundreds of nanometers to several microns from the main *Tremblaya* cell body (**Figure S2**). The morphology and scale of TLPs suggested that the *Tremblaya*-associated membrane system may form extensive interconnections with host cell organelles. Notably, TLPs frequently appeared near regions densely populated with host organelles, including mitochondria, raising the possibility that they may increase the membrane surface area for *Tremblaya*-organellar interactions, lipid exchange, or metabolic trafficking. These observations motivated us to pursue large-volume FIB-SEM imaging of an entire bacteriocyte to determine the 3D organization of the *Tremblaya*-associated membrane network, and to quantify its interaction with host organelles.

### Whole-cell reconstruction reveals widespread contacts between Tremblaya and host mitochondria and lysosomes

To determine the three-dimensional organization and interaction of *Tremblaya* with its host cell, we acquired a 4-nm isotropic resolution FIB-SEM volume (~30,826 µm^3^) of a complete bacteriocyte (**Figure 2**). To automate segmentation, we trained a deep-learning algorithm^19^ on 1285 manually annotated cutouts (**Figure 2C**), and fully reconstructed *Tremblaya, Moranella*, mitochondria, lysosomes, and the host nucleus (**Figure 3, Movie S1**). The reconstructed bacteriocyte contained six *Tremblaya* cells, 163 *Moranella* cells, 2,522 mitochondria, and 506 lysosomes. *Tremblaya* and *Moranella* cells make up 53.6% of the total bacteriocyte volume (**Figure 3E, 3F**). The 2,522 mitochondria, 506 lysosomes, and nucleus make up 6.6%, 0.9%, and 6.7% of the total bacteriocyte volume, respectively (**Figure 3B-D, Movie S1**). Combined, these annotations account for 67.8% of the bacteriocyte cell volume. A significant portion of the remaining cell volume is comprised of rough endoplasmic reticulum (RER) (**Figure S3**), but we were unable to quantify this precisely because of persistent false-negative and false-positive annotations of rough ER-trained segmentations. Using a conservative RER model, we estimated that 2.8% of the total bacteriocyte volume to be RER. We believe this to be an underestimate as regions can be seen to contain RER but failed to segment (**Figure S3B**). Our full FIB-SEM volume contained additional, partial bacteriocyte cells, and this same RER model appeared to segment RER much better in these adjacent bacteriocytes (**Figure S3C**). While these adjacent bacteriocytes are incomplete, our RER model predicted 8% and 6.1% of these adjacent cell volumes to be occupied by RER (**Figure S3D**).

**Figure 2:**
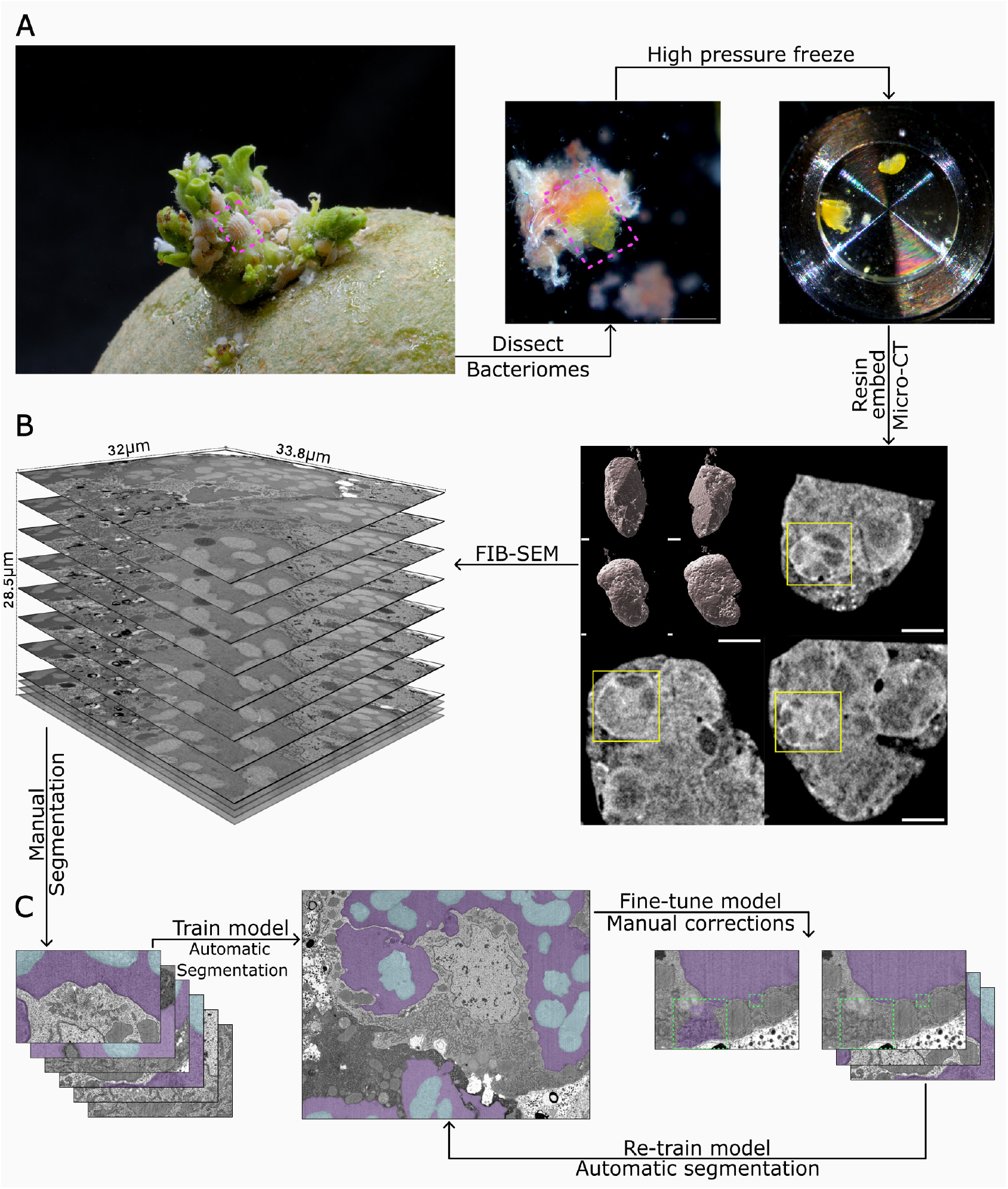
Workflow of FIB-SEM imaging and deep-learning-based segmentation of a complete P. citri bacteriocyte. (A) Overview of the sample preparation workflow. *P. citri* bugs were grown on potatoes, the bacteriomes were dissected, loaded into high pressure freezer hats and frozen, freeze substituted and resin embedded. Magenta dashed boxes indicate a *P. citri* bug feeding on a potato sprout, and subsequent dissected bacteriome. (B) Resin embedded samples were imaged via microcomputed tomography (micro-CT) to identify a well-preserved bacteriocyte for targeted FIB-SEM. The whole bacteriome is displayed from multiple viewing angles in 3D. Yellow boxes indicate the bacteriocyte selected for FIB-SEM shown in XY, XZ, and YZ views. FIB-SEM acquisition of the targeted bacteriocyte was collected at 4 nm isotropic voxel resolution over a volume measuring 32.0 × 33.8 × 28.5 µm. Representative serial images are shown. (C) Deep-learning segmentation workflow. Manually annotated image patches were used to train an initial segmentation model, which was applied to the full FIB-SEM volume for automated segmentation. Segmentation errors were identified by manual inspection. Corrected annotations were added to the training set, and the model was iteratively fine-tuned and reapplied until satisfactory segmentation accuracy was achieved. Representative segmentations show *Tremblaya* in purple, and *Moranella* in grey. Green dashed boxes show segmentation errors and corrected annotations. Scale bars: (A) 1 mm; (B) 20 µm.

**Figure 3:**
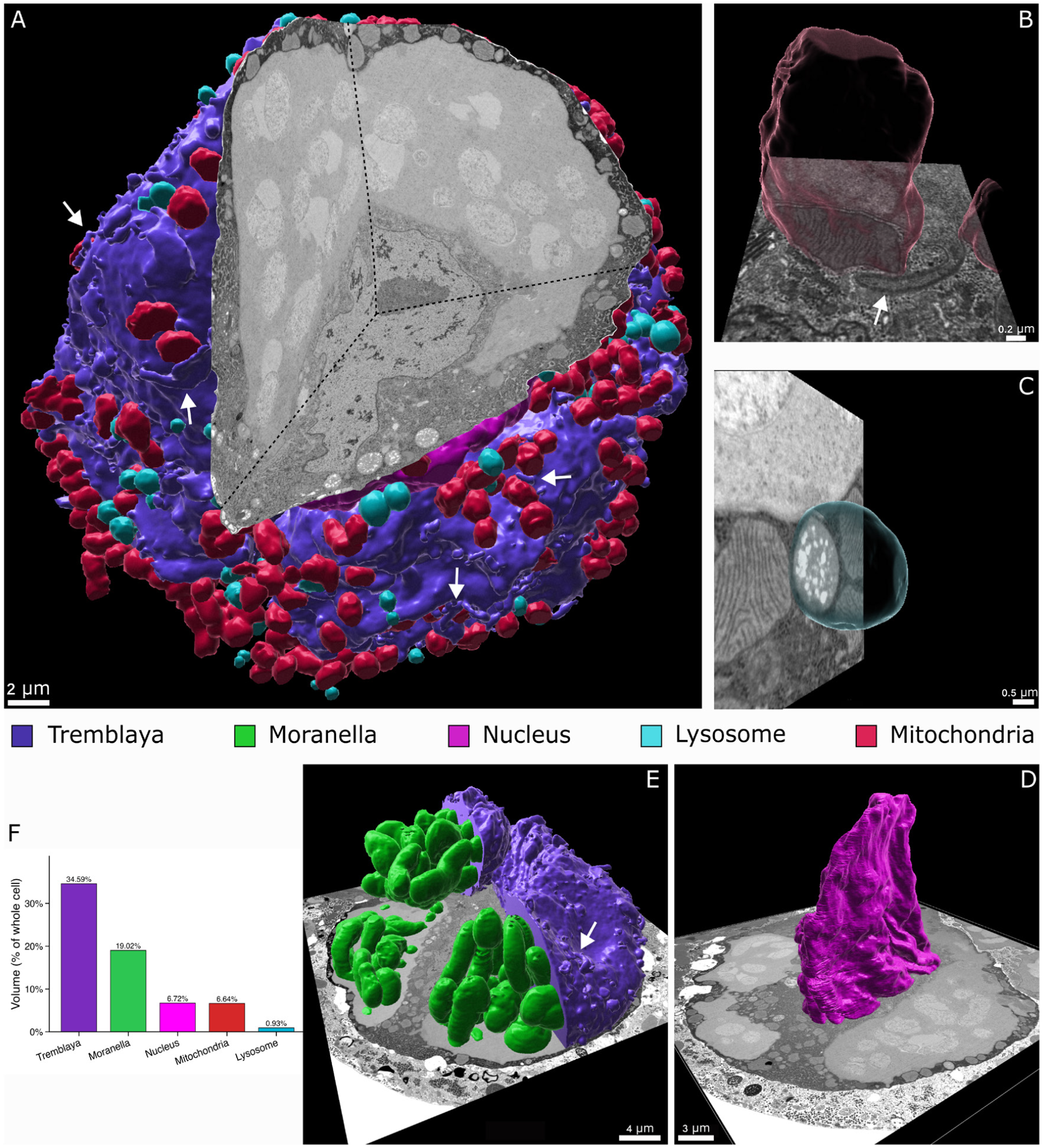
Three-dimensional reconstruction of a P. citri bacteriocyte. (A) Three-dimensional reconstruction of the segmented bacteriocyte overlaid with orthogonal FIB-SEM sections. White arrows indicate tentacle-like projections (TLP) extending from *Tremblaya* into the bacteriocyte cytoplasm, often associated with mitochondria and lysosomes. (B) Representative mitochondrion shown in close proximity to *Tremblaya* with an orthogonal FIB-SEM section. White arrow points to a TLP partially wrapping around the mitochondrion. (C) Representative lysosome shown in close proximity to *Tremblaya* with an orthogonal FIB-SEM section. (D) Three-dimensional reconstruction of the bacteriocyte nucleus overlaid on an orthogonal FIB-SEM section. (E) Three-dimensional reconstruction of *Tremblaya* and its nested symbiont partner, *Moranella*. White arrow points to a TLP. (F) Percentage of the total bacteriocyte volume occupied by *Tremblaya, Moranella*, the nucleus, mitochondria, and lysosomes. Scale bars are indicated in each panel.

Consistent with our TEM and tomography investigations, the FIB-SEM reconstruction revealed numerous TLPs throughout the host cytoplasm and extensive wrapping of individual mitochondria by *Tremblaya* cells (**Figure 3A, 3B, 3E; Figure S4; Movie S2**). All confidently identified TLPs remained connected to the *Tremblaya* cell body, although we stress that detached vesicles derived from the symbiosomal membrane are difficult to exclude here as they are nearly impossible to distinguish from host-derived vesicles using FIB-SEM methods.

Reported membrane-to-membrane distances between MCSs in other system vary, with 5-50 nm^9^, 10-80 nm^18,20^, and most commonly <30 nm^6,21–24^ being reported. For automated analysis of the full FIB-SEM dataset, we classified a mitochondrion as “contact-positive” when any portion of its surface was positioned within 30 nm of *Tremblaya*. This operational cutoff was used to provide a standardized and conservative measure of proximity and was distinct from the manual identification and measure of electron-dense bridge-associated MCSs. Using this criterion, 1,349 of 2,522 mitochondria (~53%) formed contacts with *Tremblaya* (**Figure 4**). Among contact-positive mitochondria, an average of 10.9% (±0.3% SEM) of the mitochondrial surface was positioned within 30 nm of *Tremblaya* (**Figure 4B, MovieS3)**. Contact coverage ranged from small, localized appositions to near complete mitochondrial envelopment (**Figure 4, MovieS3**).

**Figure 4:**
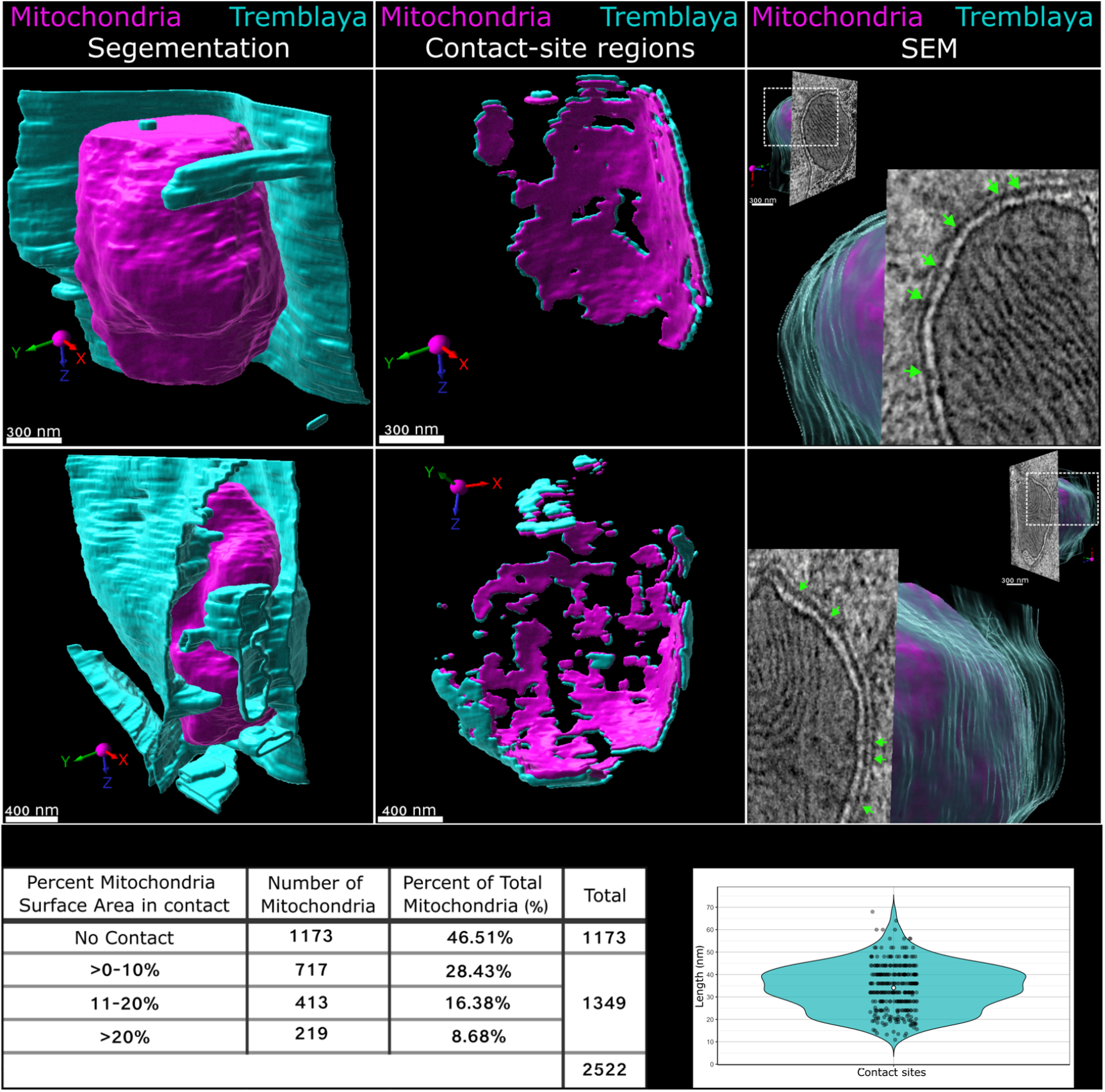
Mitochondria form extensive membrane contact sites with Tremblaya. (A) Representative three-dimensional reconstructions of mitochondria-*Tremblaya* associations. Left, complete segmentations demonstrate the extent to which *Tremblaya* membranes wrap around individual mitochondria. Middle, isolated MCS regions where mitochondria and *Tremblaya* membranes are within 30 nm of each other. Right, three-dimensional segmentations overlaid with SEM slices. Dashed white boxes show regions that were zoomed in on. Green arrows indicate regions where electron-dense bridge-like structures can be seen spanning the two apposing membranes. (B) Table show the distribution of mitochondria according to the percentage of mitochondrial surface area within 30 nm of *Tremblaya*. Of 2522 mitochondria, 1173 showed no detectable contact, 1349 formed contacts in covering >0-10%, 11-20%, or >20% of their surface area. (C) Distribution of directly measured membrane-to-membrane distances at 337 visually validated mitochondria-*Tremblaya* MCSs. MCSs spacing averaged ~34 ± 1 nm, and ranged from ~11-68 nm. Scale bars are indicated in each panel.

While analyzing the FIB-SEM dataset, we also observed contacts between host lysosomes and *Tremblaya* (**Figure S5A**). Because lysosomes form MCSs involved in lipid, ion, and metabolite exchange^7,25–30^, we applied the same analysis to the segmented lysosome population as was applied to mitochondria. By these measures, 233 of 506 (~46%) lysosomes formed contacts with *Tremblaya* (**Figure S5B**). Among contact-positive lysosomes, an average of 9.0% (±0.6% SEM) of the lysosomal surface area was positioned within 30 nm of *Tremblaya*. In several examples, the *Tremblaya*-associated membranes curved around substantial portions of the lysosome (**Figure S5A, S5B**).

We manually validated our automated bridge-associated contact estimates between *Tremblaya* and both organelles. We inspected 149 randomly chosen mitochondria predicted to have membrane contact sites by our automated method (based on a membrane-to-membrane distance of <30 nm). Of those 149 mitochondria, 142 (~95%) were confirmed to have obvious electron-dense bridges between the mitochondria and *Tremblaya* (**Figure S6**). We analyzed a subset of those 149 mitochondria (n = 47) and measured the length of 337 distinct electron-dense bridges between the outer mitochondrial membrane and the *Tremblaya*-associated symbiosomal membrane. These individual bridges had an average membrane-to-membrane distance of 34 ±1 nm and ranged from 11 to 68 nm (**Figure 4C**). For lysosomes, we inspected 36 randomly chosen lysosomes predicted to have membrane contact sites, and 30 of the 36 (~83%) were confirmed to have electron-dense bridges between lysosomes and *Tremblaya*. We analyzed a subset of those 36 lysosomes (n = 12) and measured the length of 337 distinct electron-dense bridges between the lysosomal membrane and the *Tremblaya*-associated symbiosomal membrane. These MCSs had an average membrane to membrane distance of 31 ±1 nm and ranged from 18 to 61 nm (**Figure S5C**). This manual inspection of *Tremblaya*-organelle contacts shows that our 30 nm cutoff in our automated calculation of MCS surface area is conservative and our estimate of the extent of these MCSs is likely an underestimate.

### Contact-mitochondria exhibit distinct external and internal architecture

Next, we tested whether mitochondria in contact with *Tremblaya* differed morphologically from mitochondria elsewhere in the bacteriocyte. Contacting mitochondria were significantly larger than non-contacting mitochondria, exhibiting approximately two-fold greater volume (geometric mean ratio = 2.01, p ≤ 0.0001) (**Figure S7A**) and 1.5-fold greater surface area (geometric mean ratio = 1.52, p ≤ 0.0001) (**Figure S7B**). Contact-mitochondria also exhibited significantly lower surface-area-to-volume ratios (geometric mean ratio = 0.76, p ≤ 0.0001) (**Figure S7C**) and moderately greater sphericity compared to non-contacting-mitochondria (mean = 0.71 vs 0.69, p ≤ 0.05) (**Figure S5D**), indicating that their increased size was accompanied by altered overall geometry rather than proportional scaling.

The mitochondrial complexity index (MCI) is a quantitative, unitless, three-dimensional metric that describes the shape complexity of individual mitochondria^31^. Applying MCI analyses to our data revealed values that largely overlapped between contact-and non-contacting-mitochondria (geometric mean = 2.22 vs 2.55) (**Figure S7E, S8**). However, mitochondria within the top 10% of the complexity distribution were approximately twice as frequent among non-contacting-mitochondria, suggesting that highly complex mitochondrial forms are less commonly associated with *Tremblaya* (**Figure S8**).

To test for differences in the internal structures of mitochondria, we quantified cristae architecture in a subset of contacting and non-contacting-mitochondria. Contacting mitochondria exhibited significantly greater cristae surface density (mean = 28.5 vs 21.2 µm^2^·µm^-3^, p ≤ 0.0001) (**Figure 5A, 5B**) and cristae volume fraction (mean = 0.258 vs 0.182, p ≤ 0.01) (**Figure 5A, 5C**), indicating that they contained more cristae-associated surface area and cristae volume relative to their total size. Therefore, the greater size of *Tremblaya*-contacting-mitochondria is coupled with a disproportionate increase in cristae abundance, indicating that these mitochondria differ in both overall size and internal membrane architecture.

**Figure 5:**
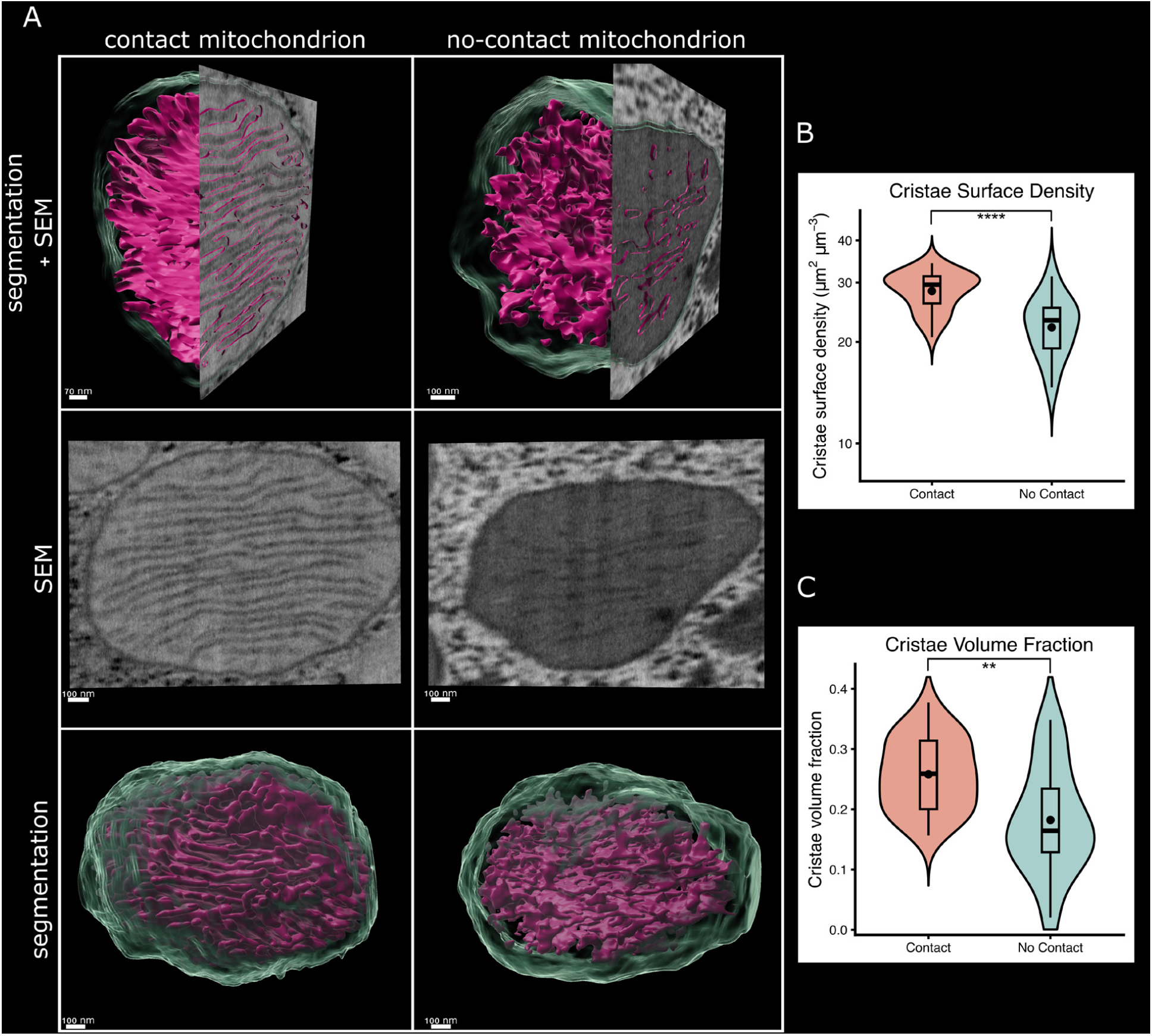
Cristae architecture differs between mitochondria that contact Tremblaya and non-contacting mitochondria. (A) Contact and non-contacting mitochondria shown as three-dimensional segmentations overlaid with orthogonal FIB-SEM slices (top row), corresponding raw two-dimensional FIB-SEM slices, and complete three-dimensional reconstructions. Mitochondrial boundaries are displayed in translucent green, and cristae in magenta. (B) Cristae surface density in contact and non-contact mitochondria. (C) Cristae volume fraction in contact and non-contact mitochondria. Violin plots show the distribution of measurements, internal boxplots indicate the median and interquartile range, and black points indicate the mean. Contact mitochondria n = 27, non-contact mitochondria n = 14. Statistical significance is indicated as: ** p ≤ 0.01, **** p ≤ 0.0001. Scale bars are displayed in each image.

## DISCUSSION

The mitochondria-*Tremblaya* contacts described here structurally resemble mitochondria-endoplasmic reticulum (ER) MCSs in other systems^32–34^. These reported ER contacts typically cover ~2-20% of the mitochondrial surface, although more extensive wrapping has been observed, and span membrane-to-membrane distances of ~25-54 nm^5,17,32,35–40^. The frequency and extent of mitochondria-*Tremblaya* contacts observed here are therefore comparable to, and in some cases exceed, those of established organelle contact sites. Together with the presence of electron-dense bridges and extensive membrane wrapping, we argue that the membrane interfaces we describe here represent specialized MCSs rather than incidental organelle crowding.

MCSs function as platforms for lipid and metabolite exchange, calcium signaling, organelle remodeling, and metabolic regulation^7,9,41^. The extensive contacts between *Tremblaya* and host organelles described here support a model where lipid acquisition, nutrient exchange, or other processes required to maintain this highly integrated symbiosis. Because *Tremblaya* lacks the genes needed to synthesize or transport its own membrane components, direct contact with host organelles provides a plausible route for acquiring molecules that it can no longer produce independently. Direct transfer across these sites remains to be demonstrated, however, and the contacts may also participate in signaling, membrane remodeling, or endosymbiont spatial organization.

Mitochondria associated with *Tremblaya* were larger, had greater surface area, lower surface area:volume ratios, moderately greater sphericity than non-contacting-mitochondria, and increased cristae density and volume. These differences suggest that contact mitochondria are not simply scaled-up versions of the broader bacteriocytes mitochondrial population but instead occupy distinct morphological and physiological states. Mitochondrial morphology is closely linked to metabolic state, fission-fusion dynamics, and organelle interactions^42–47^, and the extensive wrapping of mitochondria by *Tremblaya*-associated membranes may contribute to this remodeling. Measurements of mitochondrial shape complexity differences, however, were more subtle. Contacting and non-contacting mitochondria showed broadly overlapping distributions, but highly complex mitochondria were enriched in the non-contacting population. This may suggest that mitochondria engaged in *Tremblaya* contacts are less likely to adopt highly complex morphologies, perhaps because contact formation imposes physical or structural constraints.

That contacting mitochondria exhibit a greater cristae volume fraction and cristae surface density suggests an altered metabolic state, perhaps an increased metabolic activity, compared to non-contacting mitochondria. Cristae contain much of the oxidative phosphorylation machinery, and their organization varies with mitochondrial physiological and metabolic state^48–51^. One interpretation of these data is that mitochondria in contact with *Tremblaya* have increased energetic demands or are spatially specialized to support processes occurring at the host-symbiont interface. The increased cristae surface density could provide additional inner-membrane area for respiratory-chain complexes and ATP-synthase, possibly supporting elevated oxidative phosphorylation^52^. Cristae architecture has been linked to respiratory chain supercomplex organization^48^, and these densely packed cristae could support increased energetic and metabolic demands of lipid transfer, metabolite exchange, membrane remodeling, or other processes required to maintain this symbiotic relationship. Alternatively, cristae differences may possibly reflect mitochondrial remodeling influenced by the formation or maintenance of MCSs rather than increased respiratory activity. Mitochondrial inner-membrane structure is influenced by a variety of factors, including lipid composition, membrane curvature, fusion and fission, nutrient availability, and cellular stress^50,53–56^. Thus, the partial wrapping of mitochondria by *Tremblaya* could physically constrain or affect local signaling and lipid exchange in ways that alter cristae organization. Future studies combining ultrastructural analysis with functional measurements of mitochondrial activity, lipid composition, and metabolic flux could help tease out these possibilities.

While the presence of so many MCSs between *Tremblaya* and host organelles suggests some specific function(s), we of course do not yet know what functions these bridges provide. While we suspect lipid exchange may be of primary importance, this does not mean that *Tremblaya* membranes might only be produced by these structures. The presence of complex membrane emanations from *Tremblaya* into the host cytoplasm (**Figure 3 and S2**) also raises the possibility of vesicle exchange as a mechanism for both molecule transport between host and endosymbiont and membrane exchange through vesicle fusion. We imagine that the two processes could be linked, for example if the lipids that originate through MCSs help define the identity of these membranes for subsequent vesicle fusion with the host.

While the vast majority of bacteria retain the ability to produce their own membranes, several endosymbionts have experienced such extensive genome reduction that they are unable to produce the molecules that define their boundaries^3^. Some of these endosymbiont genomes no longer encode even a single transmembrane protein. But as their existence relies on the transport of nutrients to and from the host^57,58^ and even protein import from the host^12,59^, the origin, biochemical composition, and transmembrane protein population of these membranes is of primary concern. Our report here of extensive membrane contact sites between host organelles and a bacterial endosymbiont that lacks the ability to produce membranes suggests that specific and close host-symbiont membrane appositions may be an important part of this complex biochemical and cellular puzzle.

## MATERIALS AND METHODS

### Mealybugs

*Planococcus citri* were reared on mini pumpkins at 25C, 77% relative humidity, and 12-hour light/dark cycles in a Percival 136LL incubator until use. Second and third instar female insects were used for all imaging experiments.

### Sample preparation for TEM and FIB-SEM

Bacteriomes were fixed in 3% EM-grade glutaraldehyde and 1% EM-grade paraformaldehyde in 0.1 M cacodylate buffer for 1 h on ice. Bacteriome tissue used for FIB-SEM were incubated in 1.5% KFeCN for 2 h (KFeCN omitted for tissue used in TEM and tomography investigations), rinsed 3X in 0.1 M cacodylate buffer, and then briefly stored in 0.1 M cacodylate buffer with 10% ficoll. Bacteriomes were loaded into Type-B brass planchettes (Ted Pella, Inc.), covered with the flat side of Type-B brass planchettes and rapidly frozen with a Leica EM-Ice high pressure freezing machine (Leica Microsystems). Vitreous samples were transferred under LN_2_ to cryovials containing freeze substitution buffer (2.5% osmium tetroxide, 0.1% uranyl acetate in acetone) and moved to an AFS-2 freeze-substitution unit (Leica Microsystems). Samples were processed at −90°C for 72 h, warmed to −20°C over 12 h, held at −20°C for 12 h, then warmed to 4°Cover 2 h. The freeze substitution buffer was removed, and the samples were rinsed 3X in glass distilled acetone for 1 h per rinse, then warmed to room temperatures over 2 h. Planchettes containing bacteriomes were transferred to a Leica flat mold (reagent bath 16707154; flat embedding inserts 16707155) and infiltrated with EMbed 812 resin (Electron Microscopy Sciences) over 48 h. Resin was polymerized at 60°C for 48 h. Samples were further prepared for FIB-SEM as described elsewhere^60,61^. Embed 812 resin blocks were re-embedded in Durcupan and mounted on a copper stud. 3D X-ray tomography (Xradia Versa 730, Carl Zeiss) was used to identify bacteriocytes within the resin block with ideal morphology to maximize the milling volume. Samples were trimmed using an ultramicrotome (Leica UC7) to form a block face with dimensions of approximately 100 × 100 x 65 µm containing the bacteriocyte of interest. Samples were then coated with 10 nm Au and 100 nm C (Gatan model 682) before being mounted into a custom sample holder FIB-SEM imaging.

### TEM and Tomography

Semi-thick sections (500-1000 nm) were cut with a UC7 ultramicrotome (Leica Microsystems) using a diamond knife (Diatome, Ltd.), and then stained on a glass slide with 1% methylene blue to determine the preservation quality. Semi-thick sections were imaged on a Nikon Ti-2 confocal microscope (Nikon) using a 20X objective (0.75 NA). For traditional TEM imaging, ultra-thin sections (60-70 nm) were cut with a UC7 ultramicrotome (Leica Microsystems) using a diamond knife (Diatome, Ltd.) and placed on Formvar-coated copper slot grids (Ted Pella, Inc.). Grids were imaged with a Talos L120C transmission electron microscope (120 KeV; Thermo Fisher) equipped with a Ceta-S camera. For dual-axis tomography imaging, semi-thick sections (150-400 nm) were cut with a UC6 microtome (Leica Microsystems) using a diamond knife (Diatome, Ltd.). Sections were placed on Formvar-coated copper-rhodium slot grids (Electron Microscopy Sciences) and stained with 3% uranyl acetate and lead citrate. Ten nm gold beads were placed on both sides of the grid to serve as fiducial markers for imaging and alignment. Grids were placed in a dual-axis tomography holder (Model 2040, E.A. Fischione Instruments) and imaged with a Tecnai TF30ST-FEG transmission electron microscope (300 KeV; Thermo Fisher) equipped with a 2k x 2k CCD camera (XP1000; Gatan, Inc.). Tomographic tilt-series were acquired using the SerialEM software package^62^. For tomography, samples were tilted ± 64° and images collected at 1° intervals. The grid was then rotated 90° and a similar series taken about the orthogonal axis. Tomographic data were calculated and analyzed using the IMOD software package^63^.

### FIB-SEM

FIB-SEM imaging was performed using modified Zeiss Gemini 450 FE-SEM equipped with Capella FIB column for right angle milling, as described by Xu et al.^64^. SEM imaging was done using 0.9 kV electron landing energy and 0.5 nA current. Images with dimensions of 33.8 × 34.1 µm were acquired with 4 nm pixel resolution at 500 kHz readout rate using an InLens detector.

The block face surface was ablated using a customized Capella FIB column equipped with gallium liquid metal source operating at 30kV accelerator voltage and 15nA aperture. Actual FIB dosage on the sample was adjusted to ablate approximately 4nm thickness per imaging cycle, and maintained using a closed feedback loop as described in Xu et al.^64^ to attain highly uniform milling. Approximately 8500 images were serially acquired spanning a depth of 35 µm. Individual FIB-SEM images were registered with respect to each other using a Python implementation of the scale-invariant feature transform (SIFT) algorithm^65^ and regularized RANSAC affine transformation solver, available at https://github.com/gleb-shtengel/FIBSEM_gs_py. Image post-processing was performed in Matlab.

### Segmentation/Annotations

In order to build image models for automated segmentation, we utilized a deep-learning-driven approach using the Empanada-Napari Plugin^19^. We manually annotated patches (512 × 512 pixels each) containing the organelle (or object) of interest (along with patches containing just background) and then trained the initial models using architecture for 10,000 iterations. 3D inference (or 2D inference in batch-mode) was run to check the accuracy of the predicted segmentations. If needed, additional patches were picked and manually annotated, followed by finetuning the model and 3D inference to check for accuracy. The final models were trained with the following number of manually segmented patches: Mitochondria, 145 patches; Nucleus, 154 patches; Lysosome, 66 patches; Tremblaya/Moranella, 920 patches. 3D inferences were generated in 10-24 separate sub-volumes to reduce the memory usage. Each chunk was exported as a zarr file. These chunks were processed using a Python script to merge labels at chunk boundaries and concatenate the sub-volumes. Imaris 10.2.0 (Oxford Instruments Bitplane, Zurich, Switzerland/Abingdon, UK) was used to visualize and proofread the segmentations. Labels were loaded and overlayed with the FIB-SEM data to check for spurious segmentations, which were removed using a combination of surface morphology filters and manual editing. Imaris was also used to produce the images in Figure 3, Figure 4, Figure 4, Figure S3, Figure S4, and all supplemental movies.

### Quantification & Visualization

MATLAB was used to extract image sub-volumes containing each mitochondrion and lysosome segmentation instance. From these sub-volumes, morphological descriptors (detailed below) were calculated for each organelle. The presence of any *Tremblaya* segmentation within the same volume was then used to detect MCSs by calculating the pairwise distance between each point on the organelle and *Tremblaya* surface, respectively. Any point pairs whose distance fell below the threshold value were deemed to constitute part of a contact site. This allowed us to identify all mitochondria near *Tremblaya*, their corresponding morphology, as well as determine how many mitochondria form MCSs with *Tremblaya*, and to what extent.

### Mitochondrial complexity and contact analysis

The MATLAB regionprops3 function was used to calculate morphological descriptors for each organelle, including surface area (*SA)*, volume (V), solidity, sphericity, and mitochondrial complexity index (MCI), using the following formulation^31^:

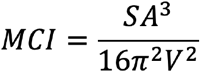

Contact between mitochondria and *Tremblaya* was defined as above. Mitochondria were classified as *contact* if any portion of their surface was within the defined distance threshold (30 nm) and *no contact* otherwise.

Because MCI values were strongly right-skewed, comparisons between groups were performed on log-transformed values using Welch’s t-test. Effect sizes were reported as geometric mean ratios, obtained by back-transforming differences in log-transformed means.

To assess robustness, additional analyses were performed. A Mann–Whitney U test was applied to untransformed data. A sensitivity analysis was performed by excluding the top 10% of MCI values. Enrichment of high-complexity mitochondria was assessed using a chi-square test, comparing the proportion of mitochondria in the top 10% of MCI values between groups. No outlier removal was performed.

### Mitochondrial cristae segmentation and quantification

To compare inner-membrane architecture between contact and no-contact mitochondria, individual mitochondrial sub-volumes were selected from the FIB-SEM dataset. Contact between mitochondria and *Tremblaya* was defined as above. Briefly, mitochondria were classified as *contact* if any portion of their surface was within the defined distance threshold (30 nm) and *no contact* otherwise. Sub-volumes contained the raw FIB-SEM data and the corresponding mitochondrial segmentation mask. Cristae were annotated in the XY, XZ, and YZ planes to train a segmentation model using the Empanada-Napari Plugin^19^. The model was applied to the mitochondrial sub-volumes to generate predicted cristae. Cristae predictions were restricted to the corresponding mitochondrial mask. Segmentations were quality control inspected so that off-target predictions outside the mitochondrion were removed. The final dataset contained 27 mitochondria and 14 non-contact mitochondria. Imaris 10.2.0 (Oxford Instruments Bitplane, Zurich, Switzerland/Abingdon, UK) was used to visualize segmentations.

For each mitochondrion, total mitochondrion volume (***V***_*mitochondrion*_), total segmented cristae volume (***V***_*cristae*_), and cristae surface area (***SA***_*cristae*_) were calculated using a python script. Cristae volume fraction was calculated as follows:

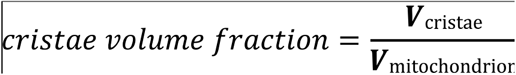

Cristae surface density was calculated as follows:

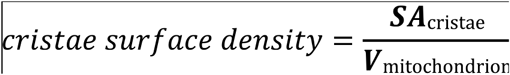

Cristae volume fraction was reported as a dimensionless proportion, and cristae surface density was reported in µm^2^/µm^3^.

Statistical analyses were performed in R-studio^66^ (2025.5.1.513). Cristae volume fraction and surface density were compared between contact and non-contact mitochondria using two-sided Wilcoxon rank-sum tests. Data were visualized as violin plots containing embedded boxplots showing the median and interquartile range; black points indicate group means.

## DATA AVAILABILITY

All data reported in this study and code used for analysis will be shared by the lead contact upon request. Raw FIB-SEM volume, segmentations, and segmentation models have been deposited to S3 and Neuroglancer: https://figshare.com/s/d74ee4533e1b3326a679?file=69266014 Raw FIB-SEM dataset used in this study have been deposited to the Electron Microscopy Public Image Archive^67^ (EMPIAR): accession code EMPIAR-13995. Supplemental movies S1, S2, and S3 are available at FigShare: 10.6084/m9.figshare.34113153.

## ACKNOWLEDGEMENTS

This study was supported by the Howard Hughes Medical Institute along with a NASA Interdisciplinary Consortia for Astrobiology Research grant (80NSSC23K1357; J.P.M) and a National Institutes of Health grant (NIH U54 AI170856; P.J.B.). We would like to thank Dewight Williams from the Eyring Materials Center at Arizona State University supported in part by NNCI-ECCS-1542160 for training us to high-pressure freeze, freeze substitute, and image on the Talos L120C. FIB-SEM imaging was performed in collaboration with the Advanced Imaging Center at Janelia Research Campus, a facility generously supported by the Howard Hughes Medical Institute. Electron tomography was performed with electron microscopes maintained by the Caltech Cryo-EM Center.

## AUTHOR CONTRIBUTIONS

Conceptualization, J.P.M.; Mealybug dissections, D.J.L; high pressure freezing and freeze substitution, D.J.L., M.S.L.; transmission electron microscopy and electron tomography imaging, M.S.L., D.J.L.; FIB-SEM imaging, J.A.; image segmentation, quantification, and visualization, D.J.L., J.A., E.H.; writing – original draft, D.J.L, J.P.M.; writing – review and editing, D.J.L, J.P.M., J.A., E.H., M.S.L., P.J.B.; Funding and oversight, J.P.M., P.J.B.

