## Supplemental Figures and Movies for "A bacterial endosymbiont with a highly reduced genome forms extensive membrane contact sites with host organelles"

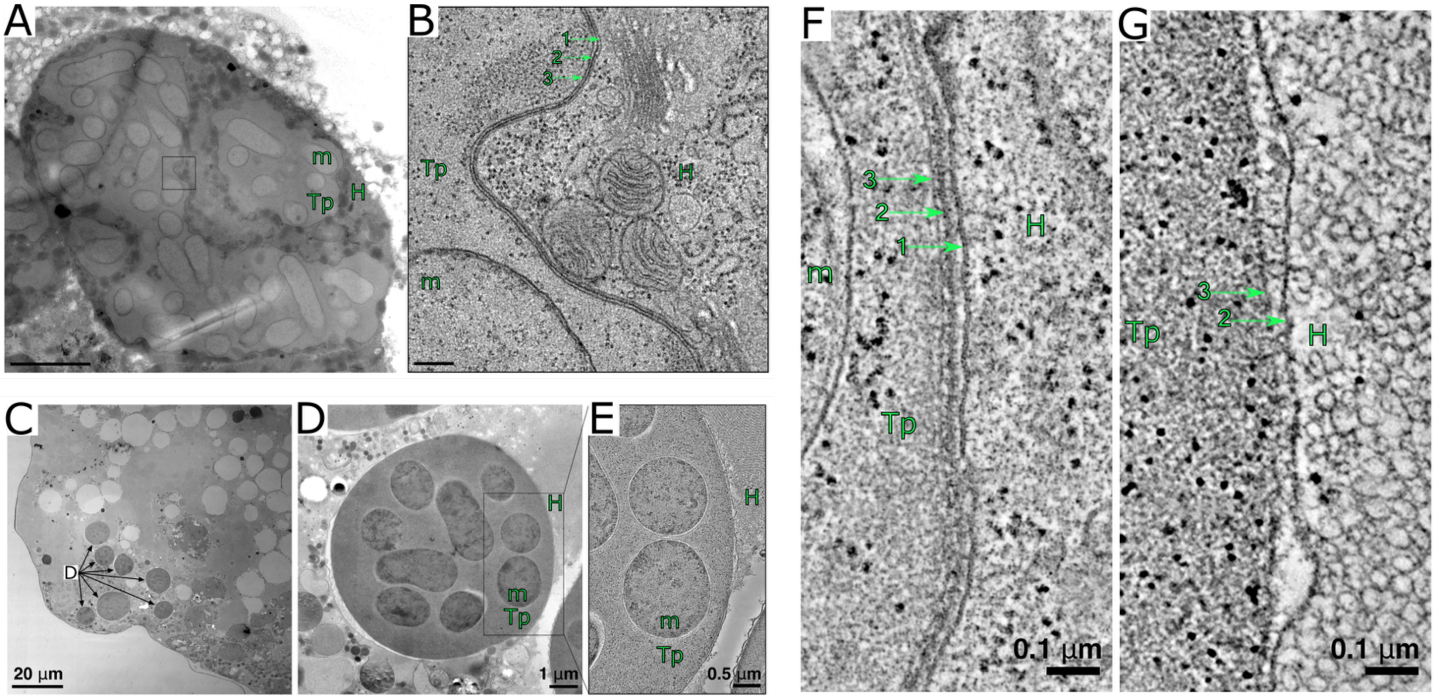


**Figure S1: *Tremblaya acquires a third membrane after entry into the host bacteriocyte*.**
(A) TEM micrograph of the nested symbiosis in *P. citri*, where *Tremblaya* resides in host bacteriocytes and contains a nested symbiont, *Moranella* in its cytoplasm. (B) Zoomed in region of the box in "A". Label 1 denotes the host-derived symbiosomal membrane, labels 2 and 3 denote the ancient didermal outer, and inner membranes of Gram-negative bacteria. (C-E) TEM micrograph of *Tremblaya* cells (arrows in D) within the symbiont ball of a *P. citri* egg prior to being phagocytized by developing bacteriocytes. (F, G) TEM micrographs of a *Tremblaya* cell in an adult bacteriocyte (F) and in the egg symbiont ball (G). *Tremblaya* is surrounded by three membranes in adult bacteriocytes, but only two membranes while in the egg. Abbreviations: Tp, *Tremblaya*; m, *Moranella*; H, host cell. Scale bars: A, 5 μm; B, 0.2 μm; all others are indicated in each panel.

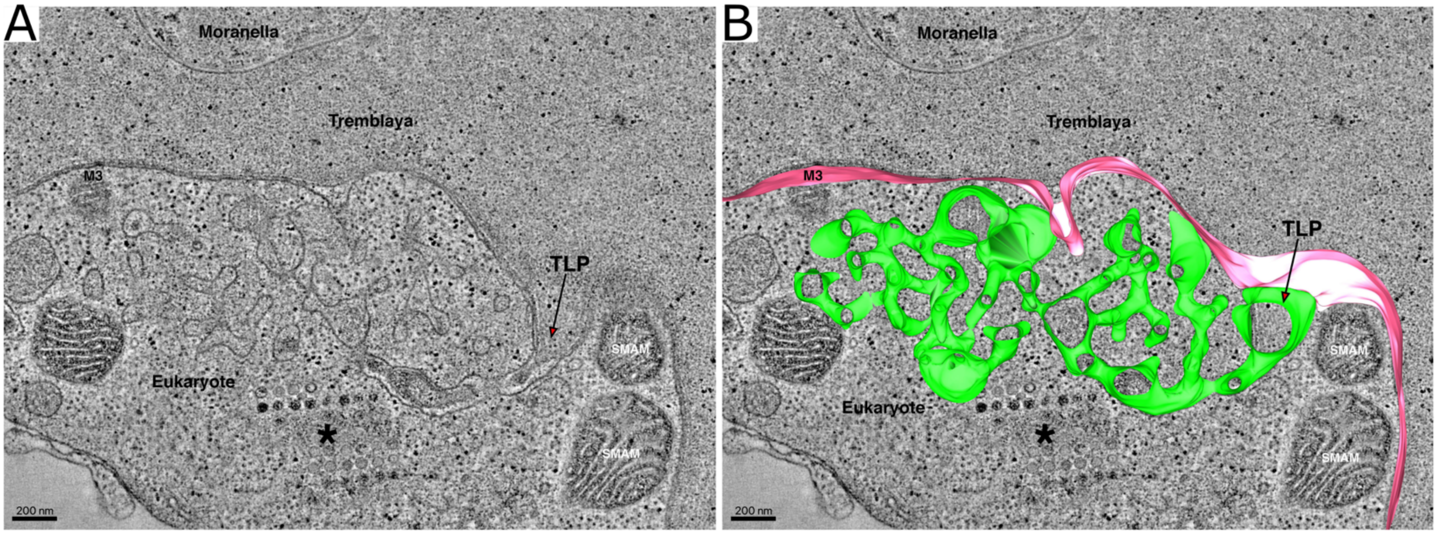


**Figure S2: *Tentacle-like projections protrude from the Tremblaya-associated symbiosomal membrane into the host cytoplasm*.**

(A) Representative tomographic slice displaying a tentacle-like projection (TLP) extending from the symbiosomal membrane surrounding *Tremblaya* into the bacteriocyte cytoplasm. TLPs are observed near mitochondria. (B) Segmentation overlay of the tomogram reconstruction. The reconstruction reveals that the TLP is continuous with the symbiosomal membrane rather than isolated vesicles in the host cytoplasm. The TLP, and diderm membranes, are displayed in green, and magenta, respectively. Scale bars: A-B, 200 nm.

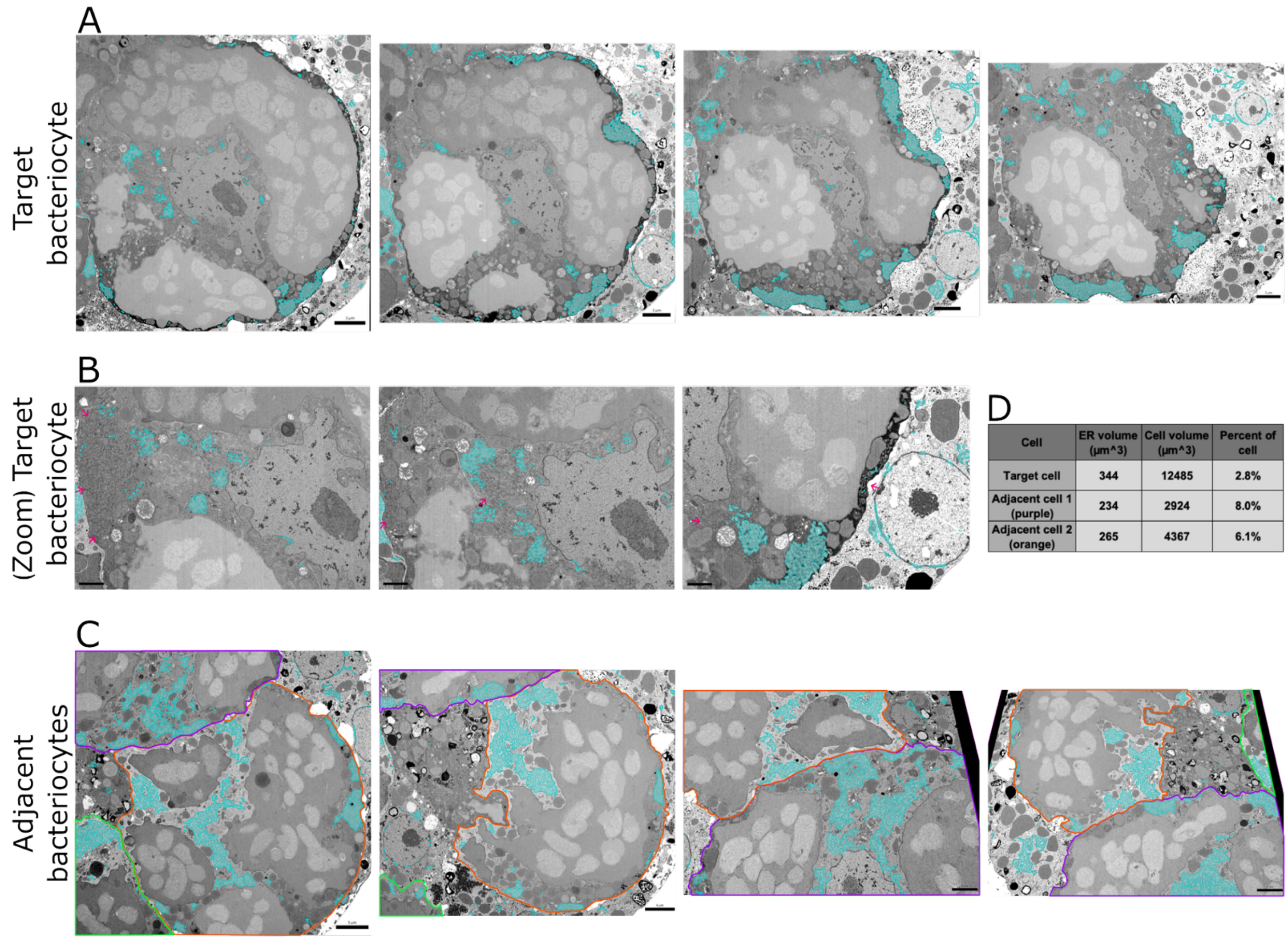


| Cell | ER volume (µm^3) | Cell volume (µm^3) | Percent of cell |
|---|---|---|---|
| Target cell | 344 | 12485 | 2.8% |
| Adjacent cell 1 (purple) | 234 | 2924 | 8.0% |
| Adjacent cell 2 (orange) | 265 | 4367 | 6.1% |

**Figure S3:** ***The endoplasmic reticulum segmentation model likely underestimates the amount of RER in the whole bacteriocyte.***
(A) Micrographs from the FIB-SEM volume of our target bacteriocyte. RER segmentations are displayed in cyan. Micrographs are shown at various z-depths. (B) Zoomed in regions from 'A.' Magenta arrows point to regions where the RER segmentation model fails to segment what is likely endoplasmic reticulum. (C) Micrographs from the FIB-SEM volume of bacteriocytes adjacent to our target cell. Cell boundaries are outlined. RER segmentations are displayed in cyan. Micrographs are shown at various z-depths. (D) Predicted RER and cellular volumes from our RER segmentation model. The target cells full volume is captured in the FIB-SEM volume. Overall bacteriocyte volumes in adjacent cells 1 and 2 are much lower than the target cell due to portions of those cells being out of the FIB-SEM volume dimensions. RER within these adjacent cells appears to segment better and occupies ~2-3X more cellular volume than the target cell. Scale bars are indicated in each panel.

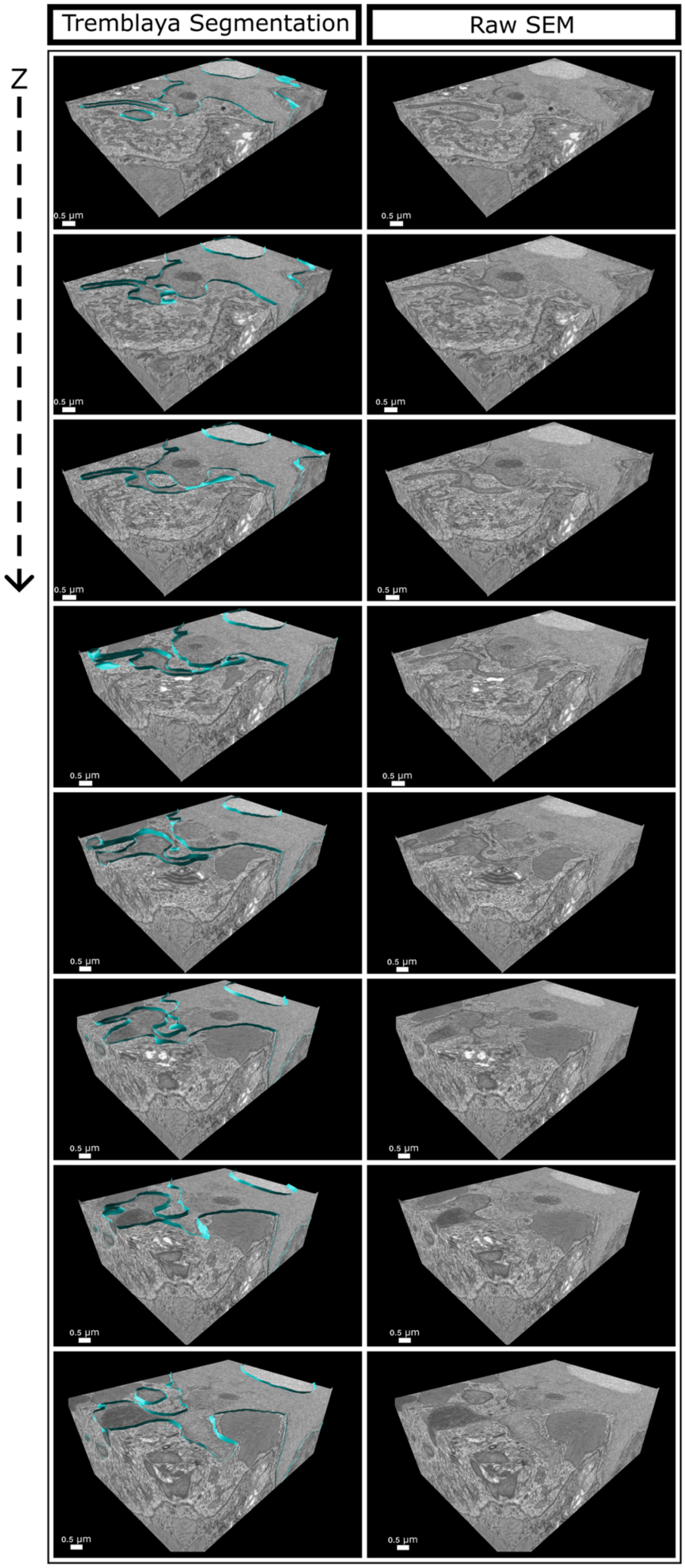


**Figure S4: *Three-dimensional continuity of Tremblaya-associated tentacle-like projections*.**
Sequential sections through a representative FIB-SEM subvolume are shown from top to bottom along the z-axis. Left, *Tremblaya* and its associated membrane projections are outlined in cyan and overlaid on the FIB-SEM data. Right, the corresponding raw FIB-SEM sections are shown without the segmentations. Tentacle-like projections (TLPs) that appear isolated from *Tremblaya* in select two-dimensional sections are revealed across successive z-planes to remain continuous with *Tremblaya.* Scale bars are indicated in each panel.

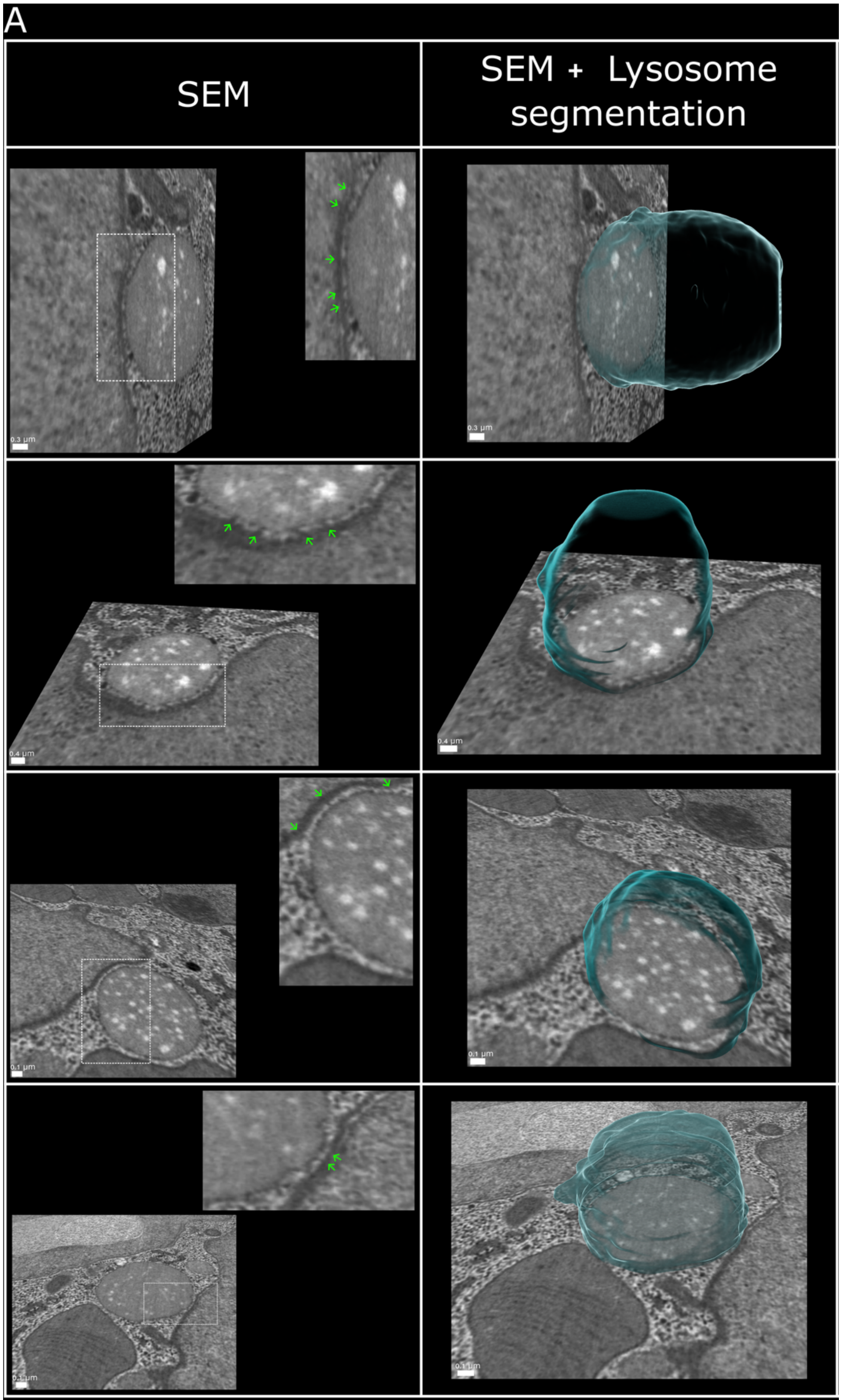


B

| Percent Lysosome Surface Area in contact | Number of Lysosomes | Percent of Total Lysosomes (%) | Total |
|---|---|---|---|
| No Contact | 273 | 53.95 | 273 |
| >0-10% | 160 | 31.62 | 233 |
| 11-20% | 42 | 8.30 | |
| >20% | 31 | 6.13 | |
| | | | 506 |

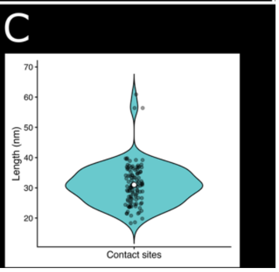


**Figure S5:** ***Lysosomes form extensive membrane contact sites with Tremblaya.***

(A) Representative FIB-SEM images of lysosomes closely apposed to *Tremblaya* membranes. Left, raw FIB-SEM section. Boxed regions are shown at higher magnification, with green arrows pointing to electron-dense, bridge-like structures spanning the space between the lysosomal and *Tremblaya* membranes. Right, corresponding three-dimensional lysosome segmentation are shown in cyan overlaid on the FIB-SEM data. (B) Distribution of lysosomes according to the percentage of lysosomal surface area positioned within 30 nm of *Tremblaya*. 273 of 506 lysosomes were further than 30 nm of *Tremblaya* membranes and classified as "no contact." 233 formed contacts involving >0-10%, 11-20%, or >20% of their surface area. (C) Distribution of directly measured membrane-to membrane distances at 116 visually validated lysosome-*Tremblaya* MCSs. Contact site spacing averaged ~31 ± 1 nm and ranged from ~18-61 nm. Scale bars are indicated in each panel.

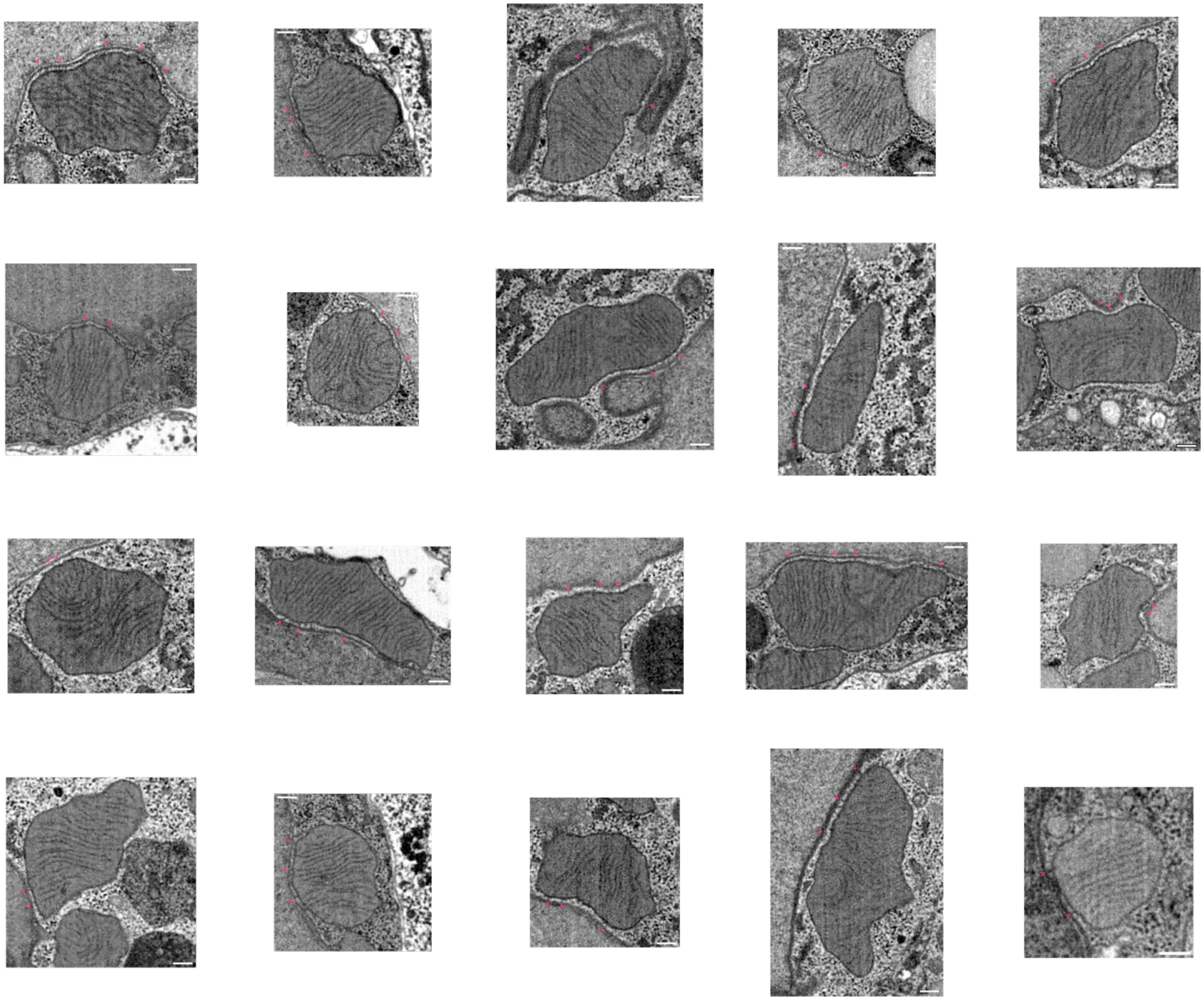

**Figure S6:** ***Examples of visually validated mitochondria-Tremblaya membrane contact sites.*** Twenty example FIB-SEM micrographs of mitochondria that were predicted by our automated analysis to form membrane contact sites with *Tremblaya* (membrane-to-membrane distance <30 nm) and were subsequently confirmed by visual inspection to contain obvious electron-dense bridges between the outer mitochondrial membrane and the *Tremblaya*-associated symbiosomal membrane. Magenta arrowheads reside in *Tremblaya* and point to representative contact regions. Scale bars = 200 nm in all panels.

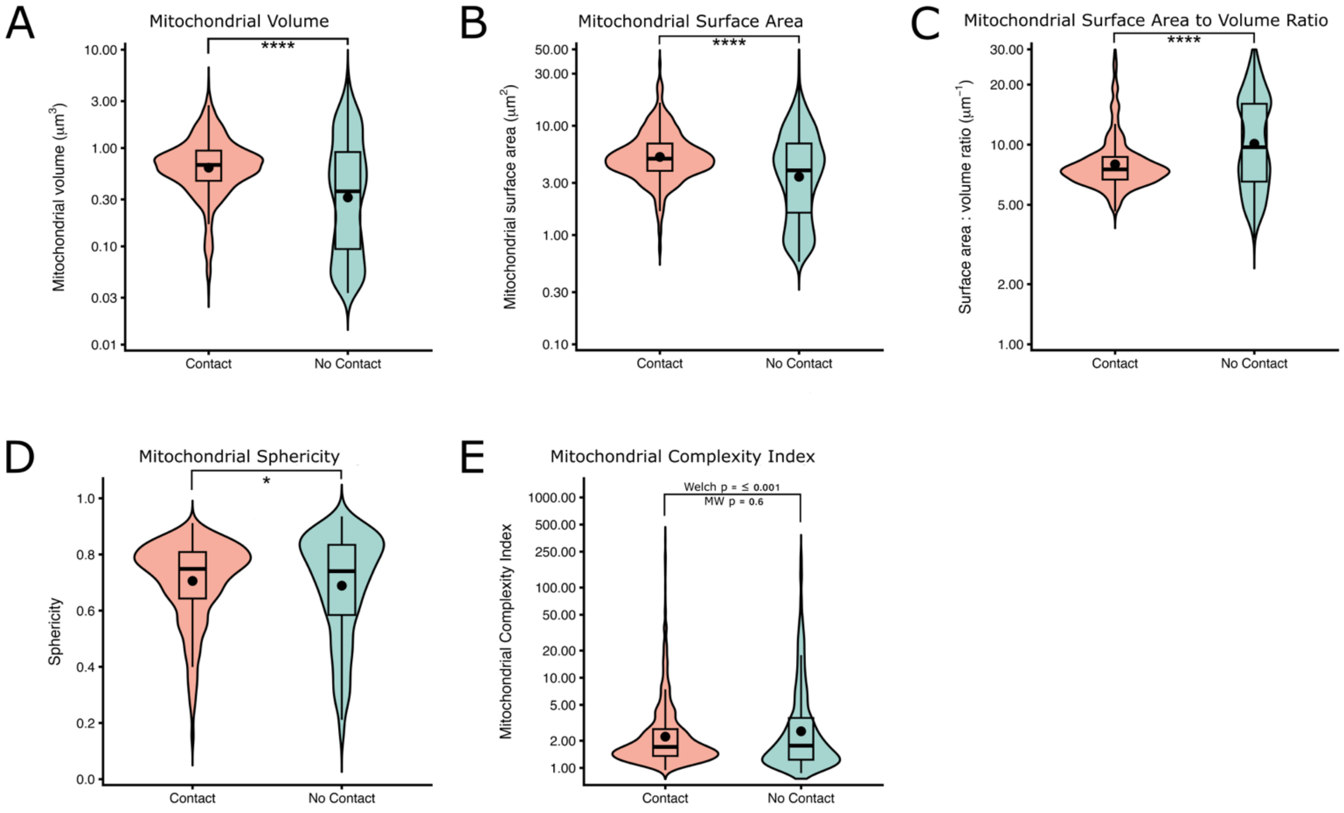


**Figure S7:** ***Morphological analyses of mitochondria associated with Tremblaya membrane contact sites.***

Comparison of mitochondrial morphology between mitochondria forming membrane contact sites with *Tremblaya* ("contact") and mitochondria not associated with *Tremblaya* ("no contact"). Violin plots show the distribution of measurements, embedded boxplots indicate median and interquartile range, and black points indicate group means. (A) Mitochondrial volume. (B) Mitochondrial surface area. (C) Surface area-to-volume (SA:V) ratio. (D) Mitochondrial sphericity. (E) Mitochondrial complexity index (MCI). Volume, surface area, SA:V ratio, and MCI were analyzed using log-transformed data and Welch's t-tests; corresponding geometric mean ratios are shown for these metrics. Sphericity was analyzed on linear-scale data. MCI distributions were highly right skewed, resulting in differing outcomes between parametric and nonparametric statistical tests (Welch's t-test $p \leq 0.001$; Mann-Whitney $p = 0.6$). Contacting mitochondria were significantly larger, exhibited greater surface area, lower SA:V ratios, and moderately increased sphericity relative to non-contacting mitochondria. Contact n = 1282; No-Contact n = 1250. $p \leq 0.05$ (*); $p \leq 0.0001$ (****).

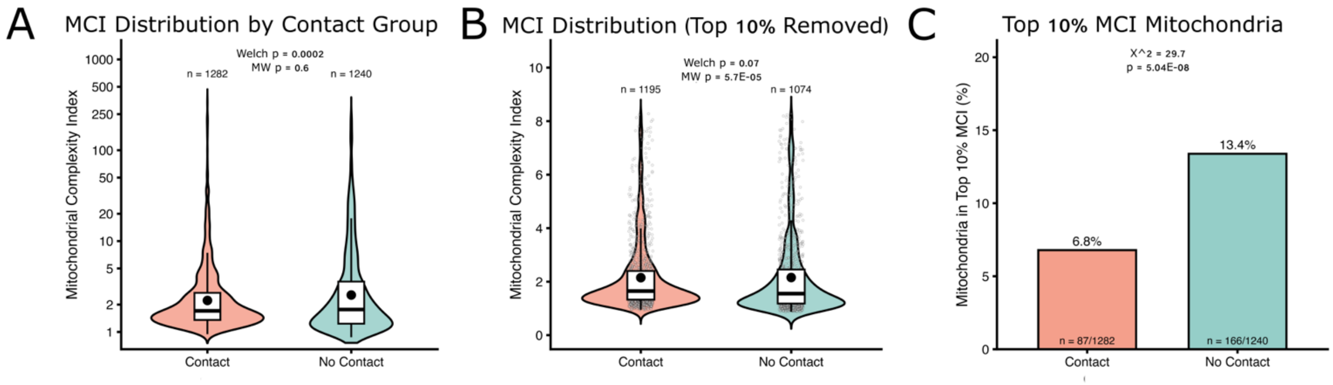


D

| Panel | Group | n | Mean MCI | Median MCI | Geometric Mean MCI | SD | SE | % Top 10% | Welch p | MW / Chi-square p | GM Ratio Contact/No Contact | 95% CI lower | 95% CI upper |
|---|---|---|---|---|---|---|---|---|---|---|---|---|---|
| A (Full MCI) | Contact | 1282 | 4.3725 | 1.7067 | 2.2206 | 16.5178 | 0.4613 | NA | 0.0002018 | 6.018e-01 | 0.8715 | 0.8106 | 0.9370 |
| A (Full MCI) | No Contact | 1240 | 6.5277 | 1.7653 | 2.5480 | 19.5072 | 0.5540 | NA | 0.0002018 | 6.018e-01 | 0.8715 | 0.8106 | 0.9370 |
| B (Top 10% removed) | Contact | 1195 | 2.1497 | 1.6489 | 1.8863 | 1.3298 | 0.0385 | NA | 0.0717900 | 5.743e-05 | 1.0395 | 0.9966 | 1.0843 |
| B (Top 10% removed) | No Contact | 1074 | 2.1551 | 1.5533 | 1.8146 | 1.5123 | 0.0461 | NA | 0.0717900 | 5.743e-05 | 1.0395 | 0.9966 | 1.0843 |
| C (Top 10% enrichment) | Contact | 87 | NA | NA | NA | NA | NA | 6.7863 | NA | 5.037e-08 | NA | NA | NA |
| C (Top 10% enrichment) | No Contact | 166 | NA | NA | NA | NA | NA | 13.3871 | NA | 5.037e-08 | NA | NA | NA |

**Figure S8: *Mitochondrial complexity index distributions differ primarily in the high-complexity tail.***

(A) Distribution of mitochondrial complexity index (MCI) values for mitochondria classified as “contact” or “no-contact” based on whether any portion of the mitochondrial surface was positioned within 30 nm of *Tremblaya*. Violin plots show the full MCI distributions on a logarithmic scale, with embedded boxplots showing the median and interquartile range. A Welch’s *t*-test on log-transformed MCI values detected a significant difference between groups ($p$ = 0.0002), whereas a Mann-Whitney U test did not detect a significant overall distribution shift ($p$ = 0.60). (B) MCI distributions after removal of the upper 10% of MCI values. Post trimming, the difference in mean log-transformed MCI was no longer significant by Welch’s *t*-test (0.07). (C) Percentage of mitochondria from each contact group represented in the upper 10% of the complete MCI distribution. High-complexity mitochondria were enriched in the non-contact group relative to the contact group (13.4% vs 6.8%; $\chi^2$ = 29.7, $p$ = 5.04 x $10^{-8}$). (D) Summary statistics for the complete MCI distributions, trimmed distributions, and the top-10% enrichment.

**Movie S1: *Three-dimensional reconstruction of a full bacteriocyte.***
Tremblaya is shown in cyan (solid and transparent), Moranella in green, nucleus in magenta, mitochondria in red, lysosomes in blue. The movie highlights the overall architecture of this bacteriocyte, and the close interactions between Tremblaya and host organelles. Scale bar is indicated in the movie.

**Movie S2: *Tremblaya tentacle-like projection wraps around host mitochondrion.***
The movie shows the three-dimensional segmentation for a portion of the *Tremblaya* cell surface (tentacle-like projection) extending into the host cytoplasm and wrapping around a host mitochondrion. *Tremblaya* is rendered in cyan, and mitochondria in red. Scale bar is indicated in the movie.

**Movie S3: *Three-dimensional reconstruction of a Tremblaya-mitochondria contact site.***
The movie first shows a FIB-SEM view of a *Tremblaya*-mitochondria contact site with obvious electron-dense bridges between the two apposing membranes. The movie then transitions into 3D reconstructions of *Tremblaya* (cyan), and mitochondria (transparent magenta). Next, it transitions to showing only regions where the two apposing membranes are within 30 nm of each other. Lastly, there is an overlay of the renderings, where transparent colors represent *Tremblaya* (cyan) and mitochondrial (magenta) membranes, and solid colors represent regions where apposing membranes are within 30 nm of each other.

*All three supplemental movies are available at FigShare: 10.6084/m9.figshare.34113153
